# Riparian Vegetation and Water Quality: A Case Study of Protected and Unprotected Rivers in the Atlantic Rainforest of Brazil

**DOI:** 10.1101/2025.04.01.646576

**Authors:** Amy Lovegrove, Eduardo Dreyer, Robin Le Breton

## Abstract

The Atlantic Rainforest, a biodiversity hotspot, has undergone significant fragmentation, with only 11% to 16% of its original cover remaining. Freshwater ecosystems within this rainforest are essential habitats, hosting high levels of biodiversity. This study assessed water quality and freshwater invertebrate diversity in rivers within and outside protected areas of the Non-Governmental Organisation Iracambi in Minas Gerais, Brazil, during the wet season from September to December 2022. We hypothesised that rivers with riparian vegetation would exhibit higher biotic indices and overall better water quality compared to unprotected rivers. Abiotic parameters air and water temperature, salinity, pH, conductivity, dissolved oxygen, flow rate, and nitrate concentration, were measured across 16 river sites, and freshwater invertebrates were surveyed to calculate Biotic Indices based on pollution tolerance. Results indicated that protected sites had a significantly higher mean biotic index compared to unprotected sites, suggesting better water quality in protected areas. Species richness was also higher in protected rivers, with 16 taxa identified compared to 11 in unprotected rivers. Pollution-tolerant species, such as *Annelida* and *Argulidae*, were more prevalent in unprotected rivers, while pollutant-intolerant species, such as *Plecoptera* and *Spinadis*, were more common in protected rivers. These findings highlight the importance of riparian vegetation in maintaining water quality and biodiversity, highlighting the need for ongoing monitoring and conservation initiatives in the Atlantic Rainforest especially in remote or isolated areas.

## 1. Introduction

The Atlantic Rainforest is the second largest rainforest in the world covering 507,900 square miles across Brazil, Argentina, and Paraguay, and is a Biodiversity Hotspot for Conservation Protection (Myers *et al*., 2000). High levels of fragmentation it has now decreased in size, with only 11 % - 16 % of original forest cover remaining in a highly fragmented ecosystem, and over 80 % of the fragments comprising less than 50 hectares (Ribeiro *et al*., 2009).

Within the forest, freshwater ecosystems are a critically important habitat for many organisms. They host high levels of biodiversity; around 9.5% of globally recognised animal species, despite comprising only 0.01% of the water on Earth (Balian *et al*., 2008). Species that are bioindicators of good water quality include mayflies (*Ephemeroptera*), stoneflies (Plecoptera), and caddisflies (Trichoptera) all of which are sensitive to pollution and require high oxygen levels to thrive, thus they are typically found in clean, well-oxygenated water (Hamid and Rawi, 2017; Tubić *et al*., 2024). Conversely, non-biting midges (*Chironomidae*), aquatic worms (*Oligochaeta*), and blackflies (*Simuliidae*) are bioindicators of poor water quality as they can tolerate much lower concentrations of oxygen in the water, and typically found in areas of high pollution and organic content (Barbosa *et al*., 2001; Oliveira and Callisto, 2010).

*Fazenda Iracambi* is a 1200-acre farmland in rural Rosário da Limeira, in the Brazilian Federal Unit of Minas Gerais. The non-governmental organisation (NGO) focuses on conserving and restoring the Atlantic rainforest in tandem with local agroforestry initiatives. The residents and staff of Iracambi and the surrounding area rely on the water provided by the rivers for daily life. From drinking water to livestock maintenance, without the natural waterways, it would not be possible to sustain the many families, communities, and visitors who reside in the municipality of Rosário de Limeira.

Dedication to conservation and restoration efforts has been in place since the NGO’s inception in 1986. The main goals of the NGO are reforestation of degraded areas, protection of local water sources used for drinking and cattle/agroforestry maintenance, and sustainability education. Minas Gerais translates to “Great Mine”, of which there are numerous focusing on the collection of iron, and bauxite for aluminium extraction. Since 2005, the intention to build aluminium mines has been strongly opposed but remains common in the state, with only one Environmental Impact Assessment (EIA) done prior to initiation.

There are several notable points of pollution into these rivers around the NGO, including household waste effluent, animals defecating close to the water bank, and pesticide runoff from horticulture. These contaminants can have a detrimental effect on the health and hygiene of both humans and animals who consume or wash with the water (Silva, 2023; Rashad *et al*., 2025). Rainfall washes these pollutants into the rivers, which means that during the wet season there could be a significant decrease in the quality of the rivers. This would be lessened by the planting of trees on the riparian zones of the rivers, as trees rely on phosphates (*e*.*g*., from pesticides) and nitrates (*e*.*g*., from manure) for growth (Mengel and Kirby, 1982), thereby preventing these compounds from entering the river systems.

Residential areas on-site continually release animal sewage onto the land that runs through the river systems, as there are no alternatives for wastewater treatment. Coffee farms, agriculture/agroforestry, and a severe fire just outside the Iracambi NGO boundaries were also point sources of pollution. There are many river tributaries that run from areas of heavy pollution into drinking water capture areas, and a previous University intern found *Coliform* bacteria in these water ways.

From an ecological perspective, higher levels of contamination in the water lead to reduced oxygen (Fu and Ono, 2023), increased turbidity (Lee *et al*., 2016), and variations in pH (Saalidong *et al*., 2022). The invertebrates and fish that inhabit the rivers cannot survive if these conditions persist, and without the river organisms, the wider freshwater ecosystem cannot function. The risk of trophic level interruptions between predator and prey, and even higher levels of turbidity from lack of filter feeder activity, increases with pollution. Therefore, the need for freshwater quality assessment is imperative so that detrimental effects can be monitored, poor-quality rivers can be restored, and good-quality rivers can be maintained.

Iracambi has hosted many research projects over the years including invertebrate diversity and water quality assessments. These have greatly contributed to understanding the previously poorly understood freshwater ecosystems of the rural are of the forest however projects are often only during the southern hemisphere winter (June to August) and efforts are not continued through the wet season (October to March). This study sought to fill this gap and provide both water quality and invertebrate species abundance and diversity during the Brazilian summer by addressing the following aims:

1. To provide comprehensive water quality and freshwater invertebrate assessments for the river system within areas protected by the NGO, and in areas outside of protection and,
2. To determine whether a significant difference exists in the biotic and abiotic measurements between the two areas.

We hypothesised that rivers with riparian vegetation will have a significantly higher biotic index than those without protection, and that overall water quality (based on several parameters) will also be higher in protected rivers.

## 2. Methods

### 2.1. Selection of river sites

Data were collected during the wet season from September 28^th^ to December 28^th^ 2022. Sites were selected based on their proximity to known pollution within the Iracambi NGO land and outside in the local countryside. There were eight rivers in open, unprotected areas and eight in areas that were well covered by riparian vegetation (Figure 1). They were chosen for their function (hygiene, plant maintenance, drinking *etc*.), location, proximity to potential pollutants, and robustness of previous data collection. Elevation, longitudinal and latitudinal coordinates were taken at each site, and the geographical range of all sites span the NGO land and into central Graminha.

**Figure 1.**
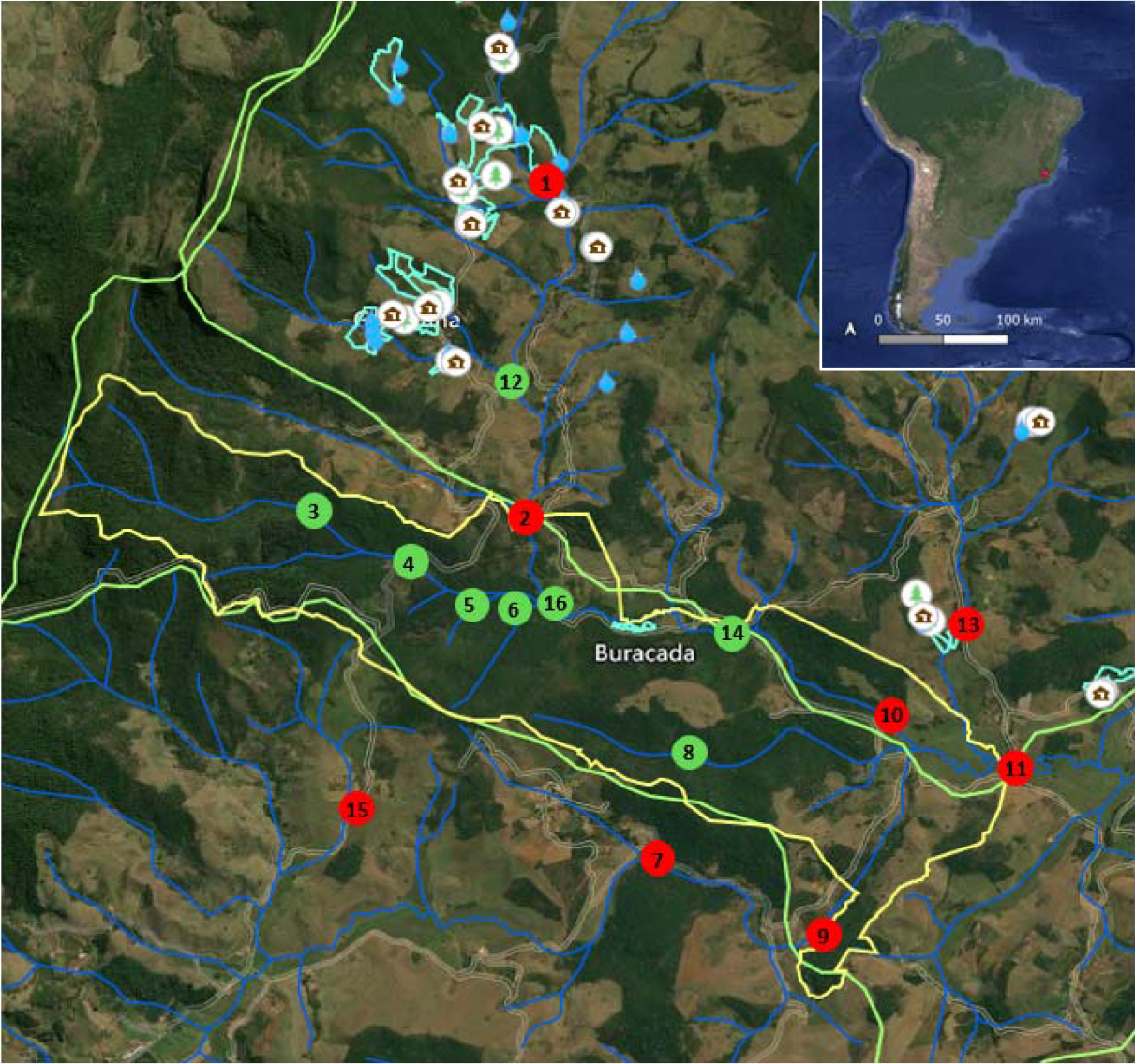
ArcGIS map of river systems within the Iracambi Fazenda and surrounding areas, and map of Brazil in top right with red dot showing where Iracambi is located. Iracambi NGO outlined in yellow, river systems shown in blue, Instituto Estadual de Florestas (IEF, State Forestry Institute) Environmental Protection Area outlined in green. The sites in red are in open, unprotected riparian zones, and sites in green are in well-protected riparian zones. The blue droplets indicate freshwater springs, the brown houses in the white circles are family farms, and the tree in a white circle represents an area of reforestation efforts. The close collection of several house and droplet icons is the village of Graminha.

### 2.2. Measurement of abiotic parameters

The methods and equipment used here were designed to be easily accessible and understandable to residents and staff who did not have any scientific training so that the water quality could be realistically continued. Air and water temperature, salinity, and flow rate were all determined in-situ using equipment outlined in Table 1, and a sample of river water was collected in a 200 mL plastic bottle.

**Table 1.** Materials list for each abiotic parameter measured. Measurement units are given for each parameter except for pH which is not applicable.

| Parameter | Units | Equipment |
| --- | --- | --- |
| Air temperature | °C | TDSTestr, Eutech, UK. |
| Water temperature | °C | TDSTestr, Eutech, UK. |
| Salinity | ppm | TDSTestr, Eutech, UK. |
| Conductivity | $\mu\text{S cm}^{-1}$ | Lutron Intelligent Meter. |
| Nitrates | ppm | LaMotte Octa-Slide Code 1100. |
| Dissolved oxygen | % | LabQuest DO<sub>2</sub> probe, multiply by 1.33 to give mg L<sup>-1</sup>. |
| Water pH | N/A | LabQuest pH probe. |
| Flow rate | m s<sup>-1</sup> | Transect, neutrally buoyant cork, timer. |
| Rainfall | mm | 1 L plastic bottle (bottleneck removed) |
| Invert (FWI) sampling | N/A | Scoop net (5 mm aperture mesh), tray, camera. |

Water from the 200 mL sample bottle was decanted into the observation pane of the Octa-Slide. The film on the reverse side makes the water appear various shades of red, depending on nitrate concentration. The scale ranges from 0 to 10 ppm with the higher concentrations appearing a deeper shade of red. To measure dissolved O_2_ concentrations, the lid was removed from the sample bottle and the probe placed directly into the water.

Air and water temperature, and salinity were measured using the same temperature and salinity probe. The probe left open outside of the water for 30 seconds to stabilise the air temperature reading. It was then placed into the river for a further 30 seconds to stabilise the water temperature and salinity readings.

River-bed width and depth was measured, and area calculated. From the GPS waypoint, 2 m was measured downstream where a second person waited. A neutrally buoyant cork was placed into the river and a stopwatch started. The time taken to reach the second mark was noted and flow rate calculated using the equation given in Table 1.

The top section of a 1 L water bottle was removed and inverted to fit inside bottom half to act as a funnel to catch rainfall. It was placed in an open area of ground and checked daily for rain. On days where rain occurred, the water was decanted into a 50 mL pot and the volume measured in mL. To calculate the rainfall in mm, the measured volume (mL) should be divided by the catchment area of the bottle and multiplied by 1000. The area of the bottle was 44.18 cm^2^.

### 2.3. Freshwater Invertebrate biodiversity survey

A modified version of the traditional method of kick sampling (Wright *et al*., 1998) was used where scoops of disturbed sediment were lifted using a sieve. Muddy substrate was sampled at the same location as the abiotic variables were measured, and once completed, contents were placed into a shallow tray for identification (photographs taken if unable to identify in the field) and returned to the river upon completion. Individuals that were not able to be identified at all were given a code based on the Family.

Biotic Index (BI) was calculated using methods outlined in Stark (1998) and Batt and Pandit (2010). Values had been given to broad taxonomic groups (not necessarily Order or Family level) and ranged from one to 10 with one being the most pollutant-tolerant species, and 10 being the most pollutant-sensitive. Regardless of how many individuals of a group were identified, each group received one BI value. Not all identified groups were allocated a BI score often because the existing research about the pollution tolerance of that group is not sufficient to provide a score. Upon completion of identification, the indices were averaged to give the BI of the river.

**Table 2.** Biotic index scores of groups of invertebrates identified in the rivers. Adapted from Stake (1998) and Batt and Panditt (2010). The scale ranges from 1 (highly pollution-tolerant) to 10 (highly pollutant-intolerant).

| **Taxon** | **Common name** | **Biotic Index (BI)** |
| --- | --- | --- |
| *Annelida* | Segmented worm | 1 |
| *Araneae* | Spiders | 5 |
| *Argulidae* | Fish lice | 2 |
| *Anura* | Frogspawn | 5 |
| *Ephemeridae* | Mayflies | 4 |
| *Spinadis* | Flat-headed mayfly | 10 |
| *Bivalvia* | Bivalves | 6 |
| *Chironomidae* | Non-biting midges | 3 |
| *Corixidae* | Water boatmen | 7 |
| *Culicidae* | Mosquitoes | 4 |
| *Dytiscidae* | Diving beetles | 5 |
| *Gerridae* | Water striders | 5 |
| *Hemiptera* | True bugs | 5 |
| *Pisidium tenuilineatum* | Fine-lined pea mussel | 6 |
| *Plecoptera* | Stoneflies | 10 |
| *Zygoptera* | Damselflies | 8 |

### 2.4. ArcGIS mapping and statistical analysis

All data were uploaded into ArcGIS® (ESRI, USA) where the base map of Iracambi land, Environmental Protection Areas (EPAs), and river systems had previously been mapped. GPX documents of sample sites were uploaded via the Garmin GPS, and abiotic and FWI survey data input into separate layers as shapefiles. A student’s unpaired *t-test* was used for all comparisons of invertebrate individuals and taxa richness, and of abiotic parameters between protected sites and unprotected sites. A point-biserial correlation test was used to show how each species correlated to each abiotic parameter, and to each protection status. All statistics were performed using accessible packages in RStudio (version 4.2.2.).

## 3. Results

### 3.1. Average daily and monthly rainfall

The average daily rainfall was 9.30 mm (± 1.75 mm SEM) (Figure 2), with monthly averages for September and October at 3.58 mm (± 2.19 mm SEM), and for November and December at 14.49 mm (± 4.20 mm SEM).

**Figure 2.**
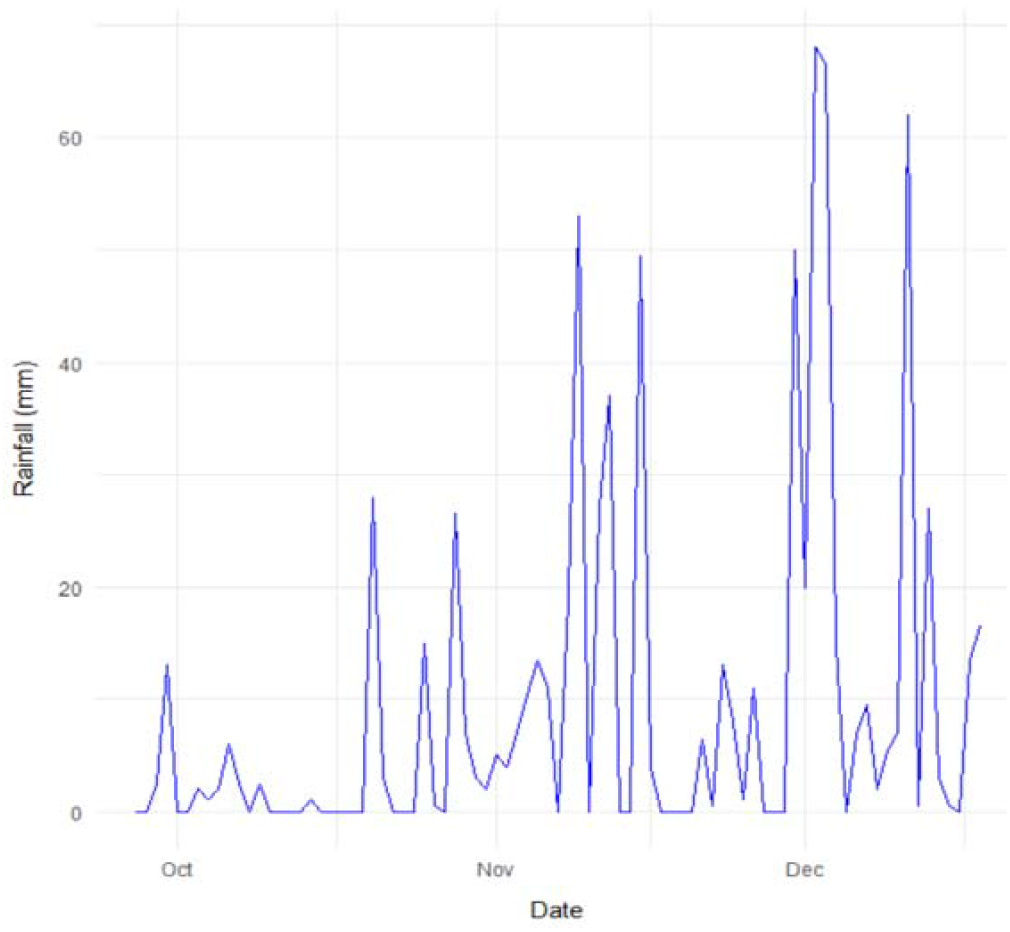
Daily rainfall (mm) from September 28^th^ to December 18^th^, 2022.

### 3.2. Abiotic water quality of rivers with and without riparian protection

For the protected sites, the average air temperature was 26.28°C (± 0.71 SEM), water temperature was 21.18°C (± 0.62 SEM), salinity was 10.63 ppm (± 0.63 SEM), and pH was 5.96 (± 0.40 SEM). The average conductivity was 21.51 µS cm^-1^ (± 2.53 SEM), dissolved oxygen (DO_2_) was 33.76% (± 1.08 SEM), flow rate was 0.19 m s^-1^ (± 0.105 SEM), and nitrate concentration was 0.031 ppm (± 0.03125 SEM).

For the unprotected sites, the average air temperature was 26.44°C (± 0.88 SEM), water temperature was 22.98°C (± 0.86 SEM), salinity was 11.25 ppm (± 2.27 SEM), and pH was 5.18 (± 0.15 SEM). The average conductivity was 20.94 µS cm^-1^ (± 2.18 SEM), DO_2_ was 29.56% (± 1.34 SEM), flow rate was 0.240 m s^-1^ (± 0.086 SEM), and nitrate concentration was 0.063 ppm (± 0.0625 SEM). The average species abundance was 43.63 (± 14.92 SEM), and average species diversity was 3.50 (± 0.60 SEM) (Figure 3). There were no significant differences between averages of the abiotic parameters at the protected versus unprotected sites (air temperature P = 0.8876, water temperature P = 0.1127, conductivity P = 0.866, flow rate P = 0.7414, pH P = 0.09691, salinity P = 0.797, nitrate P = 0.664).

**Figure 3.**
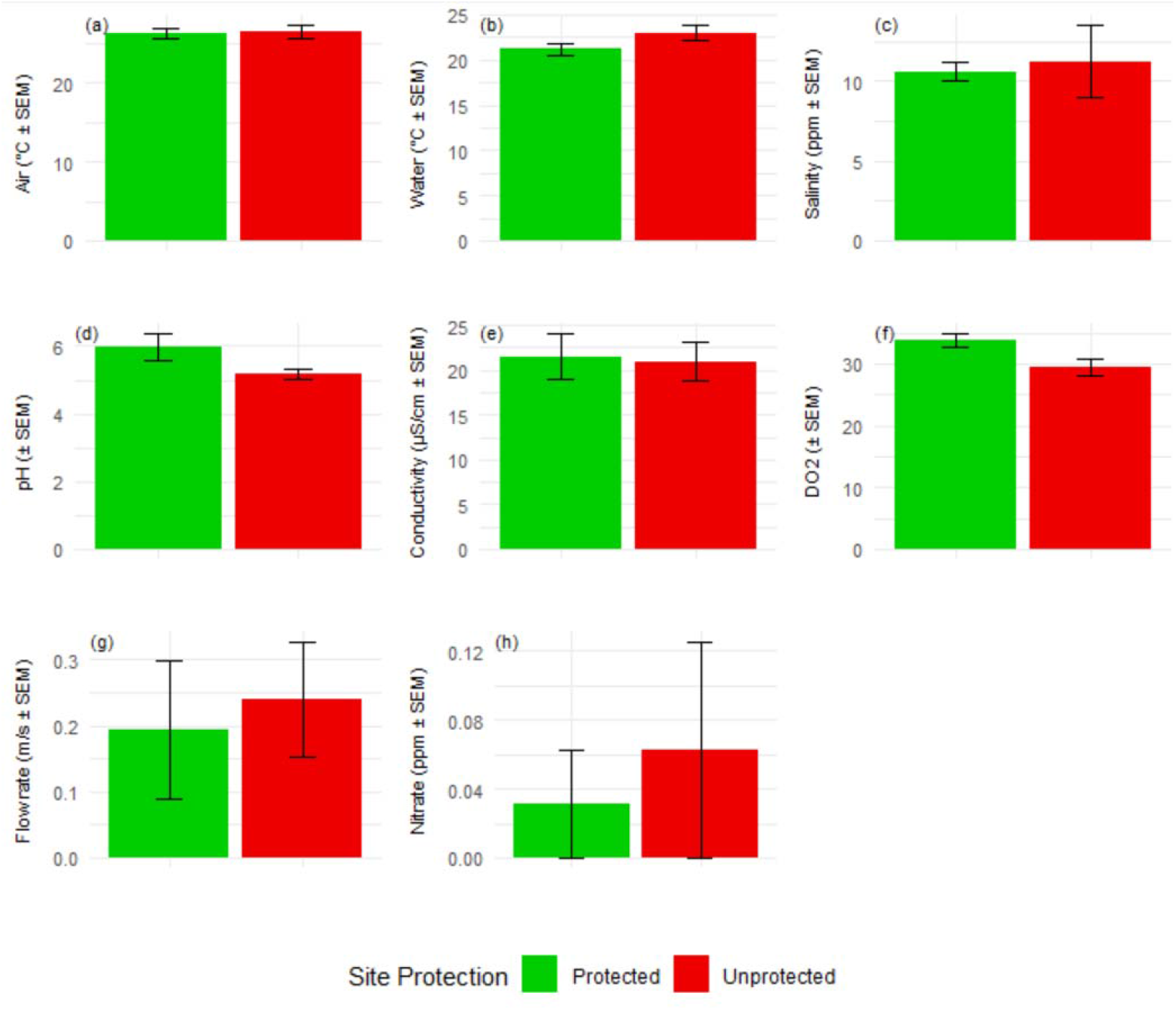
Abiotic variables for river water quality assessment for a) air temperature (° C), b) water temperature (° C), c) salinity (ppm), d) pH, e) conductivity (µS cm^-1^), f) dissolved oxygen (DO2) (%), g) flow rate (m s^-1^), and h) nitrate (ppm). All error bars are SEM.

### 3.3. Freshwater invertebrate survey

At the protected sites, the average biotic index was 6.04 (± 0.53 SEM), average abundance was 32.63 (± 11.61 SEM), and average species diversity was 3.88 (± 0.44 SEM). At the unprotected sites the average biotic index was 3.74 (± 0.41 SEM), average species abundance was 43.63 (± 14.92 SEM), and average species diversity was 3.50 (± 0.60 SEM) (Figure 4). There were no significant differences in the number of individuals (P = 0.5704) and number of species (P = 0.6221) between protected and unprotected sites. However, there was a significant difference in the biotic index (P = 0.0044), with protected sites having a higher mean biotic index compared to unprotected sites (Figure 4).

**Figure 4.**
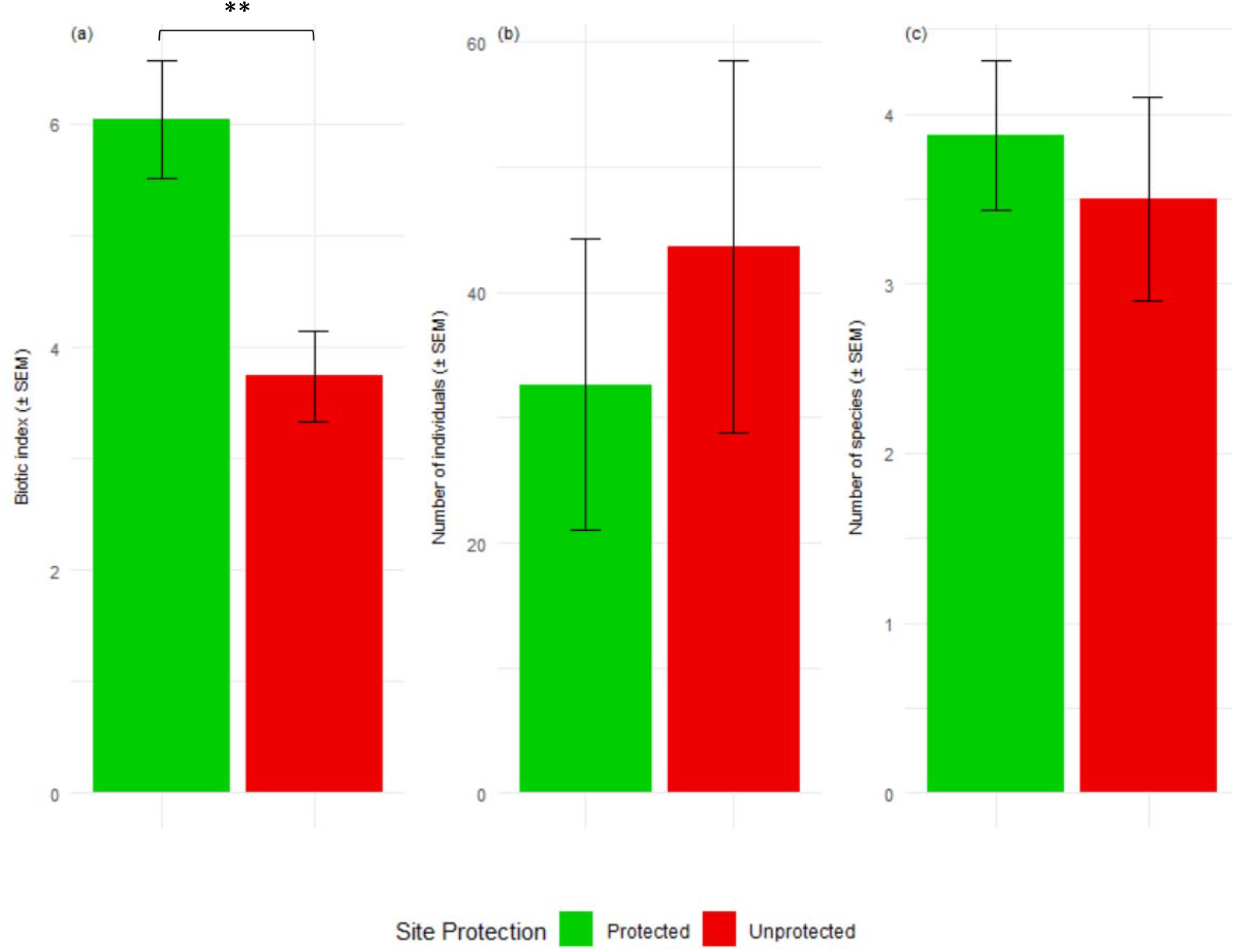
a) Biotic indices for each site, where the minimum value indicating heavily polluted water is 0 and the maximum value indicating pure, unpolluted water is 10 (*t-test P* = 0.0044**), b) average total number of individual invertebrates identified, and c) species richness of sites protected by riparian vegetation (green), and unprotected (red). All graphs include SEM.

In terms of the richness of species found in the rivers, unprotected rivers (Figure 5a) had a total of 11 different taxonomic families, including those that could not be identified grouped together and the most abundant being the water striders (family *Gerridae*) which comprise 89.01 % of the total abundance, followed by fish lice (*Argulidae*) (2.25 %), arthopods of the family *Symphyla* (1.69 %), and juvenile Anurans (1.41 %). There were 16 different taxa identified in the protected rivers (Figure 5b) again with *Gerridae* as the most abundant (52.06 %), followed by the true bugs (*Hemiptera*) (12.73 %), isopods (9.36 %), stoneflies (*Plecoptera*) (7.12 %), and the water boatmen (*Corixidae*) (6.74 %). There were five taxa in common between the two protection statuses: *Anura, Gerridae, Isopoda, Argulidae*, and *Aranae* (water spiders).

**Figure 5.**
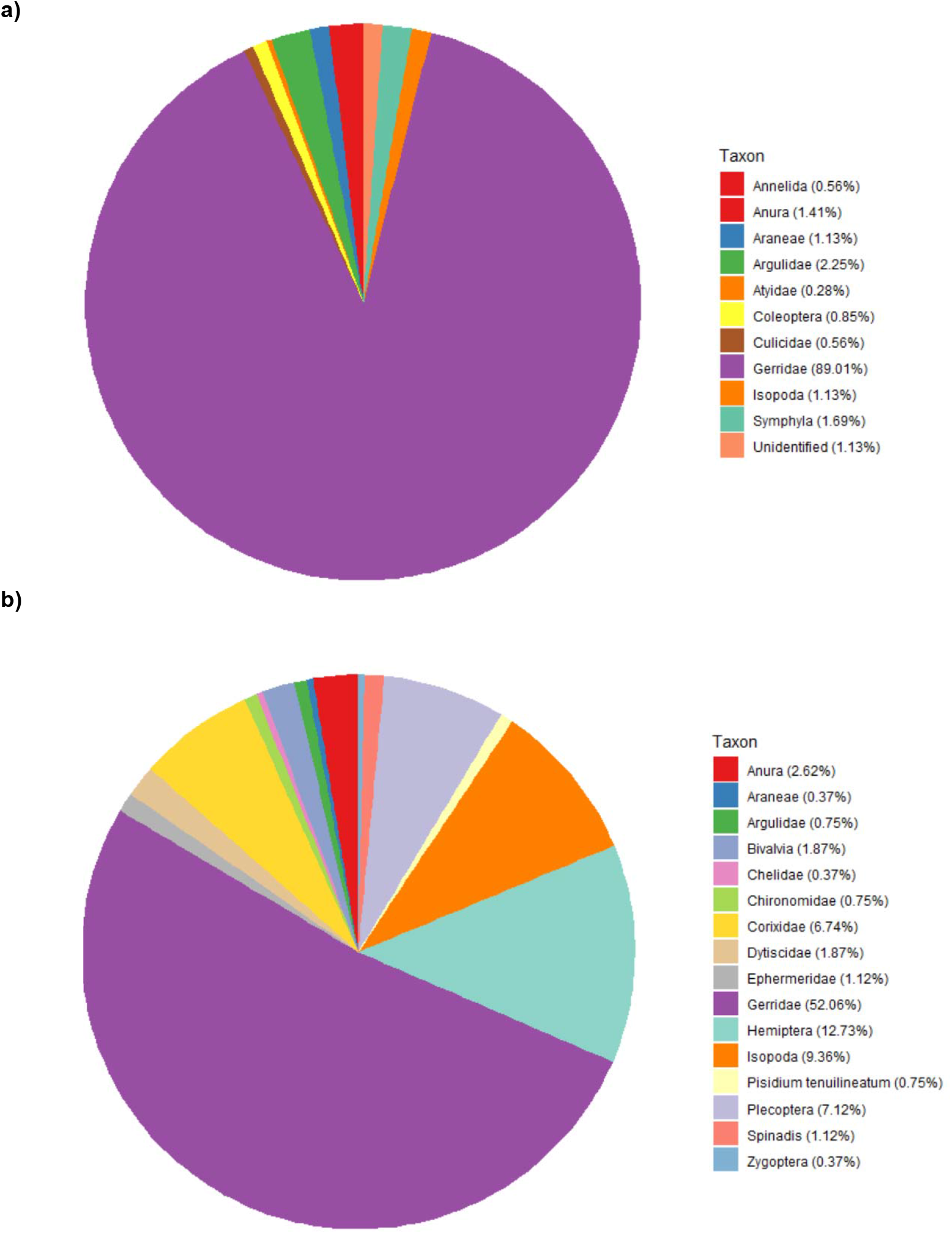
Pie charts showing invertebrate species richness in a) sites without riparian vegetation and b) sites with riparian vegetation. Percentages next to each taxon name indicate percentage of total abundance.

Based on the point-biserial correlation analysis (Figure 6), the invertebrates more likely to be found in rivers without riparian protection include *Annelida, Araneae, Argulidae, Coleoptera, Culicidae, Gerridae, Atyidae*, unidentified indivduals, and *Symphyla*. On the other hand, species more likely to be found in protected rivers include *Anura, Ephermeridae, Spinadis, Bivalvia, Chelidae, Chironomidae, Corixidae, Dytiscidae, Hemiptera, Pisidium tenuilineatum, Plecoptera, Isopoda*, and *Zygoptera*.

**Figure 6.**
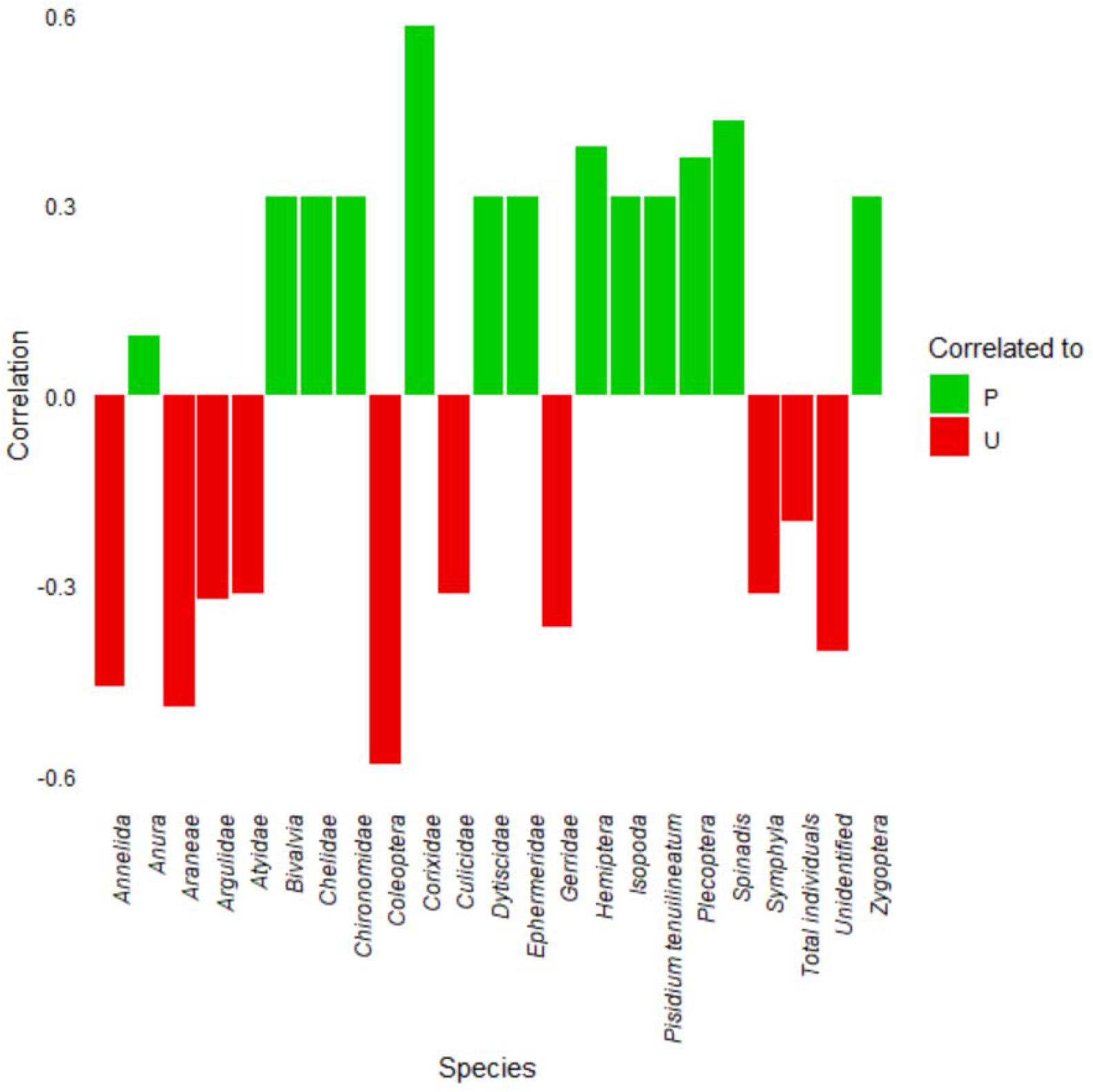
Point-biserial correlation of invertebrates identified with protected (P, green) or unprotected (U, red) rivers.

## 4. Discussion

### 4.1. The current condition of water quality in the sampled rivers

Here, we aimed to elucidate the relationship between the presence of riparian vegetation and the water quality of 16 rivers in a small rural community in the Brazilian Atlantic rainforest. This study was conducted during the wet season, where the total rainfall for the sampling period was 771.5 mm which is consistent with expected rainfall (Forti *et al*., 2005). The Water Quality Index (WQI) has shown that some parameters like turbidity, total dissolved solids, and nitrite have exceeded recommended guideline values in certain area (Avigliano and Schenone, 2016), requiring dynamic assessments of individual areas considering the local contributions to the water quality, rather than applying country-specific models like the WQI.

All abiotic parameters measured here were within the expected ranges for this time of year in the Atlantic rainforest. The average air temperatures are within the typical range for the Atlantic rainforest during this period, which experiences warm temperatures between 20 °C and 30 °C, as are the average water temperatures, which maintain temperatures between 20 °C and 25 °C. The mean pH values of 5.96 (protected) and 5.18 (unprotected) are slightly acidic, which is typical for the region, as the pH of water bodies in the rainforest often ranges from 4.5 to 6.5 due to the high organic matter content and natural acidity of the soil (Cerqueira *et al*., 2015). Conductivity typically ranges from 10 to 50 µS cm^-1^ and salinity levels are generally low, often below 20 ppm. The mean nitrate values are within acceptable limits, as in natural freshwater systems are typically low, often below 1 ppm (Dantas *et al*., 2025).

While not statistically significant, there was great variation in water quality across the rivers. Rivers with little to no riparian protection and those near pollution sources had the worst quality in terms of higher nitrate and lower dissolved oxygen, notably sites 9, 10, and 11, followed by 1, 5, 14, and 16. This difference in quality is highlighted by the preference of pollution-intolerant invertebrates (e.g., *Anura, Ephermeridae*, and *Bivalvia*) being significantly correlated to presence in protected rivers. In contrast, pollution-tolerant invertebrates (e.g., *Annelida, Araneae*, and *Coleoptera*) were significantly correlated to presence in the unprotected rivers, demonstrating their resilience to degraded conditions. Continued monitoring of the invertebrate diversity of the rivers can help guide conservation and management efforts to improve water quality and protect biodiversity in river ecosystems.

Improving all rivers to a good standard requires minimal effort, with sites 9 and 10 recommended for tree planting projects at Iracambi. Planting two rows of trees along river edges can manage pollution without encroaching on farmland, allowing cattle to drink while absorbing harmful phosphates and nitrates. Sites 1, 5, 14, and 16 need analysis during the dry season to assess turbidity changes from wet season rainfall, while repairing leaks at rivers 1, 5, and 12 is essential due to active point source pollutants from residences and animal shelters. Priority for future water quality monitoring should be given to sites 4, 8, 11, and 12, as site 4 is the main drinking source for Iracambi NGO, site 8 is closest to pristine forest and so could act as an indicator of broader environmental changes, site 11 is the most heavily polluted, and site 12 links the community and NGO.

### 4.2. The broader context of this study in the Atlantic Rainforest

Degradation of riparian zones is known to negatively impact water quality, with a recent review finding that riparian vegetation was directly related to water quality in 91% of the studies (Ebling and Padial, 2024). The impact of land use on water quality in the Atlantic Rainforest is significant, with areas having intact riparian vegetation showing better water quality and higher biodiversity compared to those affected by human activity (Avigliano and Schenone, 2016). This aligns with the findings of this study that protected rivers with riparian vegetation have higher biotic indices and overall better water quality. Agricultural runoff further exacerbates the situation, as rivers near agricultural areas exhibit higher nitrate levels and lower dissolved oxygen, leading to poorer water quality and reduced biodiversity (Muniz and Oliveira-Filho, 2023). Observations of higher nitrate concentrations and lower biotic indices in unprotected rivers support these findings. Freshwater invertebrates serve as bioindicators of water quality, with pollution-tolerant species more prevalent in degraded habitats and pollutant-intolerant species thriving in protected areas with good water quality (Avigliano and Schenone, 2016). This is consistent with results showing a higher prevalence of pollution-tolerant species in unprotected rivers and pollutant-intolerant species in protected rivers.

The importance of riparian vegetation in maintaining water quality and biodiversity is a recurring theme in the literature. Studies consistently show that areas with intact riparian vegetation have better water quality and higher biodiversity, emphasising the need for conservation efforts focused on riparian zones (Avigliano and Schenone, 2016; Muniz and Oliveira-Filho, 2023).

Fragmentation of the Atlantic rainforest has profound effects on freshwater ecosystems that can be measured using various abiotic parameters such as temperature, pH, conductivity, dissolved oxygen, and nitrate concentration as was done in the present study. The higher biotic index and species richness in protected rivers are consistent with findings from other research, suggesting that protected areas generally maintain better water quality and support higher biodiversity. For instance, a study on the Fish-based BI in the rainforest found that streams with higher biological integrity, indicated by diverse fish assemblages, were predominantly located in protected areas (Cetra and Ferreira, 2016). Additionally, research on the effects of human activities on rivers in protected areas of the Atlantic forest showed that upstream anthropogenic activities can significantly alter the natural condition of rivers, but protected areas help mitigate these impacts by maintaining better water quality and supporting aquatic biota (Kuhlmann *et al*., 2014). These findings stress the importance of conservation efforts in riparian zones to preserve the ecological integrity of freshwater ecosystems in the Atlantic rainforest.

### 4.3. Continuing monitoring efforts to protect the rivers into the future

To improve the understanding of these river ecosystems and their biodiversity, several steps are recommended for future research. Repeating the study during the dry season (April to October) would provide valuable comparative data, allowing for a seasonal analysis of river quality and biodiversity. Collaborating with residents and students to monitor environmental changes and collect data could offer continuous and comprehensive insights into the river ecosystems. Assessing the impact of newly planted trees near the riverbanks would help determine the effectiveness of reforestation efforts on river health and biodiversity.

Consistent sampling effort must be maintained to understand the implications of climate change-driven rainfall increase and anthropogenically induced ecosystem breakdown. Climate change is likely to contribute to sediment erosion in the wet season with higher levels of rain. Anecdotally, the second incident in recorded history of hail in the area was experienced during this research trip (the other occurring in November 2015). Protecting rivers with riparian vegetation is a priority, as it not only preserves biodiversity but also ensures the sustainability and resilience of these ecosystems for future generations.

By examining water quality and freshwater invertebrate diversity during a period of high rainfall, this research fills a gap in understanding seasonal variations in river ecosystems. The findings contribute to the broader understanding of the role of riparian vegetation in maintaining water quality and biodiversity, offering valuable insights for conservation efforts in fragmented landscapes which are further fragmenting as agroforestry proliferates in the forest. The results of this study have significant implications for practical management strategies in the Atlantic Rainforest. The higher biotic indices and better water quality observed in protected rivers underscore the importance of riparian vegetation in mitigating pollution and supporting biodiversity. These findings can inform conservation policies and reforestation efforts, guiding the implementation of riparian buffers to improve water quality and ecosystem health. By linking scientific research to actionable management practices, this study offers a valuable resource for policymakers and conservationists.

The biotic indices and other indicators used in this study provide a robust framework for assessing water quality and biodiversity in freshwater ecosystems. These indicators can be adapted for broader environmental monitoring programs, offering a standardised approach to evaluate the health of isolated river systems. By elaborating on the development and application of these indicators, this research contributes to the advancement of ecological monitoring methodologies, supporting the integration of scientific findings into practical conservation efforts.

## Acknowledgements

The authors would like to thank the University of Southampton SoCoBio Doctoral Training Partnership and the BBSRC for funding this work as part of the DTP placement, and the staff at Iracambi NGO who helped in navigating and sharing knowledge of the local area.

